# Elastic mitotic tethers remain functional after microtubules are inhibited

**DOI:** 10.1101/2024.05.18.594830

**Authors:** Aisha Adil, Maral Janan, Arthur Forer

## Abstract

During normal anaphase in animal cells, elastic tethers connect partner telomeres of segregating chromosomes and exert backward (anti-poleward) forces on those chromosomes. The experiments reported herein test whether microtubules need to be present in order for tethers to produce backwards forces. We disassembled spindle microtubules by treating anaphase crane-fly primary spermatocytes separately with nocodazole, colcemid, or podophyllotoxin. The drug treatments caused anaphase chromosomes to stop moving poleward; almost immediately thereafter they moved backward. The characteristics of the backward movements of the half-bivalents match those of the backwards movements of arm fragments formed by cutting chromosome arms during anaphase – for example the occurrence and lengths of backward movements were a function of tether length. The only difference from movement of arm fragments is that the chromosomes in the treated cells moved backwards slower than arm fragments did. Immunofluorescence of spindle tubulin after the drug treatments indicated that acetylated kinetochore microtubules were not depolymerized by the drugs, though the non-kinetochore spindle microtubules were depolymerized. Our data indicate that tethers move anaphase chromosomes backwards in the absence of functioning spindle microtubules. We suggest that the backward movements that take place when poleward forces are absent are due to tethers, and that the backward movements are slowed by the presence of acetylated kinetochore microtubules.

## INTRODUCTION

This study investigates whether inhibiting microtubules during anaphase in crane-fly spermatocytes affects the backward forces on chromosomes produced by elastic mitotic tethers. Mitotic tethers, present in a broad range of animal cells (Forer *et al.,* 2017), physically connect telomeres of each segregating pair of half-bivalents during anaphase in crane-fly primary spermatocytes (Forer and Otsuka, 2023). Each pair is connected in two of the four arms (LaFountain *et al*., 2002; Sheykhani et al, 2017). The presence of tethers was demonstrated by cutting (with a laser) a trailing chromosomal arm in early anaphase; the resultant arm fragment moved backward in the anti-poleward direction, across the equator, and reached its partner telomere in the opposite half spindle (LaFountain *et al*., 2002). Such backward movement is due to elastic tethers and not microtubules because it occurs only when both partner telomeres are intact (LaFountain *et al.,* 2002; Forer *et al.,* 2021); occurs for only the arm fragment piece that has the kinetochore when the arm fragment itself is cut in half (LaFountain *et al.,* 2002); and occurs when taxol stabilizes microtubule activity (Forer *et al.,* 2018). The elastic tethers connecting separating telomeres have been identified electron microscopically (Forer and Otsuka, 2023) and are quite different from microtubules. The backward movement is not caused by ultra-fine DNA bridges because DNA-bridges are not elastic, are found only in a small number of anaphase chromosomes, mostly at the centromeric regions rather than at telomeres, and when present retard or stop anaphase segregation (Chan *et al.,* 2007; Barefield and Karlseder, 2012; Gemble *et al*., 2015; Su *et al*., 2016).

Tether elasticity is diminished and then lost as anaphase progresses. When arm fragments of segregating anaphase chromosomes are cut in mid anaphase rather than early anaphase, their backward movements are less frequent than in early anaphase and the fragments move shorter distances, not reaching the partner telomeres; when formed yet later in anaphase the arm fragments do not move at all. Thus, tethers become less elastic as anaphase progresses and eventually are inelastic (LaFountain *et al.,* 2002; Forer *et al.,* 2021). Loss of tether elasticity during anaphase is due to dephosphorylation. By treating early anaphase cells with Calyculin A, an inhibitor of both serine-threonine protein phosphatase-1 and protein phosphatase 2A, tethers remain elastic: CalA added to cells in early anaphase causes entire chromosomes to move backward (telomere towards telomere) at the end of anaphase, after they have reached the spindle poles (Fabian *et al*., 2007a; Kite and Forer, 2020). These movements are because of tethers, as demonstrated using laser micro-irradiations (Forer *et al.,* 2021).

Partially lysing cells during anaphase stops poleward chromosome movements produced by spindle forces but the anti-poleward tether forces remain active and cause the segregating half-bivalents to move backward (Adil and Forer, 2024). The characteristics of the chromosome backward movements in partially-lysed cells are the same as those of arm-fragment movements (Forer et al., 2021) in most respects; for example the chromosomes move backwards more frequently and for longer distances when tethers are shorter than when tethers are longer, just as arm fragments do (Adil and Forer, 2024). The movements of arm fragments are different from chromosome movements in lysed cells in only one respect: arm fragments move faster than chromosomes. We attributed this difference to kinetochore microtubules: partial-lysis of anaphase cells does not disassemble kinetochore microtubules (Adil and Forer, 2024). Since the kinetochore microtubules remain attached to the backward moving chromosomes, we suggested that the attached kinetochore microtubules slow the movements compared to arm fragments that are not attached to anything that might slow them down. We tested this suggestion in the present article.

We asked two main questions. Are tethers still active in living cells in which spindle microtubules are disassembled? If so, does disassembling kinetochore microtubules cause faster backward movements of chromosomes, increasing speeds to those of chromosomal arm fragments? In the experiments reported herein we treated anaphase cells with various microtubule depolymerising agents in order to disrupt microtubule dynamics and to disassemble spindle microtubules. Then we measured the frequency and extent of backward chromosomal movements, and we measured backward chromosomal velocities. We used immunofluorescence to test whether spindle microtubules were disassembled. We used three microtubule- depolymerising agents. *Nocodazole* promotes microtubule depolymerization by binding and aggregating free tubulin (Jordan *et al.,* 1992). *Colcemid* promotes MT depolymerization by binding the microtubules at their plus end, thereby inhibiting their polymerization (Bryan, 1974; Yang *et al.,* 2010). *Podophyllotoxin* promotes microtubule depolymerization by binding and blocking the colchicine binding sites on the tubulin subunits, which inhibits polymerization of microtubules and speeds up their depolymerization rates (Bryan, 1974; Jordan *et al*., 1992; Hamel, 2003; Jordan and Wilson, 2004; Yang *et al.,* 2010). The results from the three inhibitors were similar. Since each acts with different mechanisms, the results are due to microtubule inhibition and not due to side effects of the drugs.

## MATERIALS AND METHODS

### Living cell preparations

Living cells of crane flies (*Nephrotoma suturalis* Loew) were prepared essentially as described by Forer (1982), and by Forer and Pickett-Heaps (1998). Briefly, testes of IV-instar larvae were dissected under a drop of halocarbon oil to prevent their dehydration; they then were washed in insect Ringer’s solution (0.13 M NaCl, 5mM KCl, 1.5 mM CaCl_2_, 3mM phosphate buffer, pH 6.8). Individual washed testes were spread out on a coverslip in a 2.5 µL drop of insect Ringers solution containing fibrinogen (10mg/mL). Cells were then fixed in place by adding 2.5 µL thrombin to the fibrinogen to form a fibrin clot (Forer and Pickett-Heaps, 2005). The clot-embedded cells on the coverslip were inverted over a drop of Ringers solution in a perfusion chamber (Forer and Pickett-Heaps, 2005) and the coverslip edges were sealed with a molten mixture of 1:1:1 vaseline, lanolin and paraffin. Ringers solution was perfused through the chamber to cover the clot-held cells, and the perfusion chamber was placed on a microscope stage to observe the living cells.

### Adding experimental agent

The cells in the perfusion chamber were observed using a phase-contrast microscope. Once the cells were at the required stage of division they were perfused according to the treatment group they belonged to. Control cells were treated with Ringers solution. Experimental cells were treated with various concentrations of one of three different microtubule inhibitors; nocodazole, colcemid, or podophyllotoxin. Inhibitor stock solutions prepared in DMSO and stored frozen were thawed and diluted in Ringers solution to make final working concentrations of the experimental drugs. Stocks of 60- and 33-mM *nocodazole* were used to make final working concentrations of 20, 30, 60 and 90 µM. Stocks of 67.3 mM *colcemid* were used to make final working concentrations of 100 and 200 µM colcemid. Stocks of 34.5 mM *podophyllotoxin* were used to make final working concentrations of 10, 20, 30, 50 and 60 µM podophyllotoxin. Of all the final working concentrations of experimental agents used, the one having highest DMSO concentration contained 0.55% (v/v) DMSO, below the 1% DMSO concentration found to have no effect on normal anaphase-I segregation [LaFountain, 1985].

Some of the nocodazole-treated cells were used for studying microtubules via immunofluorescence. Fifteen to twenty-five minutes after adding nocodazole, cells were perfused with lysis buffer and processed for staining with antibodies, as described below.

### Microscopy and data analysis

We studied the living cells using a Nikon 100X, 1.25 NA phase-contrast oil-immersion objective lens. Real-time video images were recorded on DVDs and later converted into time-lapse video sequences (.avi files) using freeware VirtualDub2. Measurements of chromosomal positions were made from individual frames using an in-house program WinImage (Wong and Forer, 2003). Chromosomal movement graphs were plotted using commercial software SlideWrite Plus 7.0.

### Fluorescent staining and confocal microscopy

The fluorescent staining procedure was adapted from Fabian et al. (2005). Briefly, control cell preparations and nocodazole-treated cell preparations were placed for 15 minutes in lysis buffer (which is essentially a cytoskeleton-stabilising buffer plus detergents) after which they were fixed in 0.25% glutaraldehyde in phosphate-buffered saline (PBS) for 2-3 minutes. Then the preparations were rinsed twice in PBS, placed in 0.05M glycine for 10 minutes (to neutralize unbound aldehyde groups), rinsed four times in PBS, and stored in PBS-glycerol 1:1 (v/v) at 4°C until used for staining. For immunostaining, stored coverslips were first washed in PBS to remove the glycerol, then rinsed briefly with 0.1% Triton X-100 in PBS, and then mixed with antibodies. All the cells were double-stained as follows. First they were treated with primary antibodies YL1/2 against tyrosinated α-tubulin (Abcam), diluted 1:200, followed by secondary mouse-absorbed Alexa 488-conjugated goat anti-rat antibody (Molecular Probes) diluted 1:50. Then primary mouse monoclonal antibody 6-11B-1 (Millipore Sigma) diluted 1:50 followed by secondary rat-absorbed Alexa 568-conjugated donkey anti-mouse antibody (Invitrogen) diluted 1:200. All antibodies were diluted in PBS, and cells were incubated in each antibody for 1 hour in the dark. At the end of each antibody incubation cells were rinsed twice in PBS, and then rinsed with 0.1% Triton X-100 in PBS (to “wet” the preparation to aid the next solution to spread across the entire slide). After the last antibody staining, coverslips were rinsed twice in PBS, and then placed in PBS-glycerol 1:1 (v/v) for 2-3 minutes. Finally, coverslips were mounted in Mowiol solution containing paraphenylene diamine (PPD) antifading agent (described in Fabian and Forer, 2005), and left in the dark for 24-48 hours to allow the Mowiol to harden. Once it hardened, coverslips with stained cells were stored at 4°C until studied using confocal microscopy. Cells were studied using an LSM 700 Zeiss Observer confocal microscope, with a Zeiss Plan-Apochromat 63X 1.4 NA oil-immersion objective lens. Images were collected using ZEN Black software, and further processed using FIJI-Image J freeware.

## RESULTS

### Control cells

Primary crane-fly spermatocytes have three pairs of bivalent autosomes and two univalent sex chromosomes. During metaphase, the bivalent autosomes and the univalent sex chromosomes line up at the equator. At anaphase the three bivalents disjoin into six half- bivalents (3 sets of partner half-bivalents), the partners of which move in opposite directions towards their respective poles (Figure 1). While the autosomes are segregating, the spindle length remains constant and both sex chromosomes remain stationary at the equator (Forer, 1966; Forer, 1980; Forer *et al.,* 2013). Once the autosomal partner half-bivalents get close to their respective poles (which takes about 20-30 minutes after the start of anaphase), the spindle starts elongating, the sex chromosomes start segregating, and the cleavage furrow ingresses (Figure 1).

**Figure 1:**
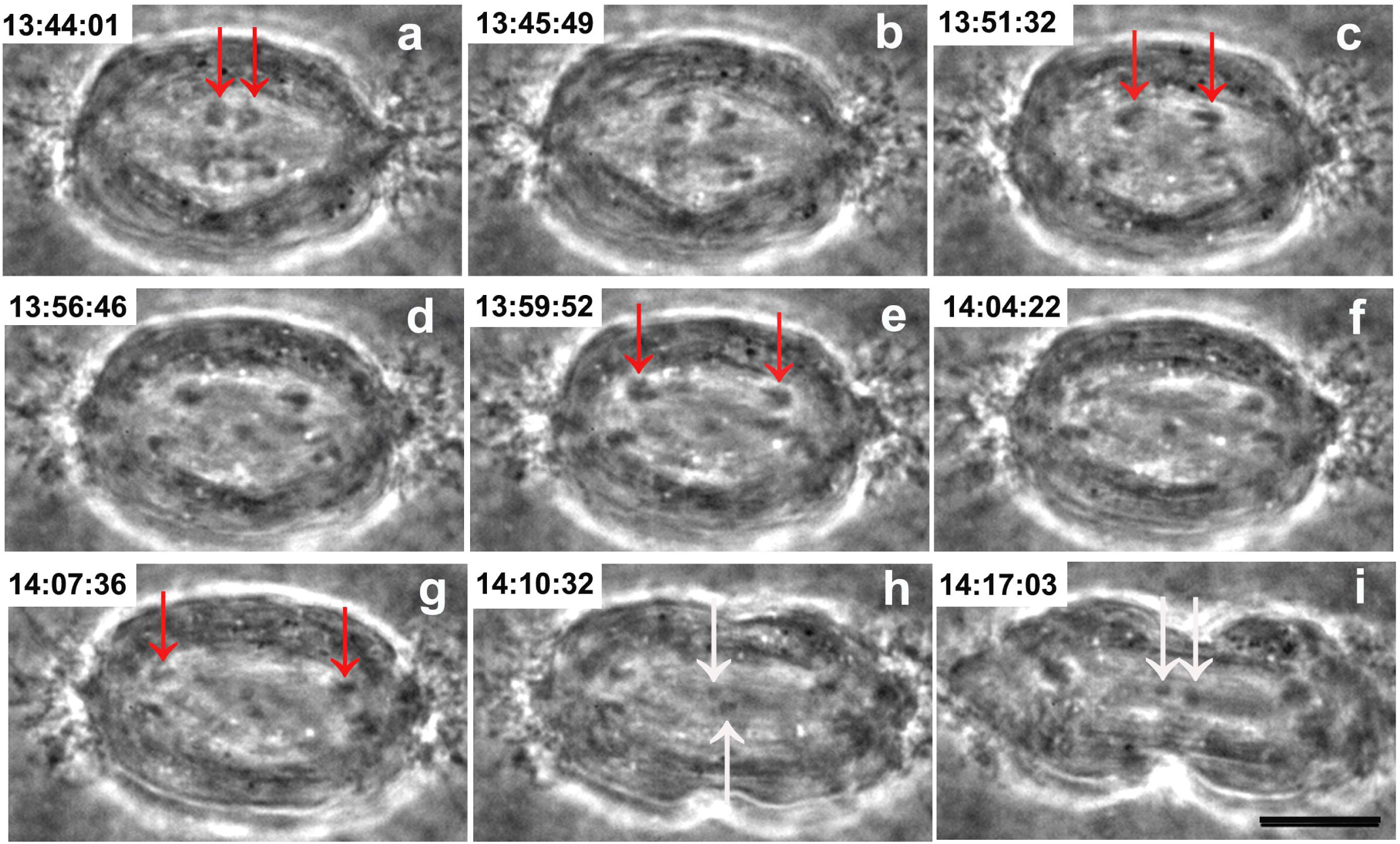
Meiosis in a control meiosis-I spermatocyte. Time in hrs:min:sec is at the top of each panel. The red arrows point to two separating autosomal half-bivalents; the white arrows point to the two separating sex chromosome univalents that move to the poles only after the autosomes have reached the poles. The cleavage furrow forms and ingresses as the univalents move poleward (**images *h-i***), and the spindle elongates as the univalents move poleward. The scale bar in ***panel i*** represents 10µm.

### Experimental cells

#### Chromosomal poleward movements stop when anaphase cells are treated with microtubule depolymerizing agents

Half-bivalents stop moving poleward and begin to move backward after microtubule depolymerizing agents are added to anaphase cells. Images of cells and graphs of kinetochore and telomere positions *versus* time are presented in Figures 2, 3 and 4 to illustrate this conclusion for each of the three depolymerizing agents. The distances between kinetochores and poles tell us whether poleward anaphase chromosome movement was stopped and whether kinetochores (chromosomes) moved backward after drug addition. The distances between partner telomeres (tether lengths) tell us the lengths of the tethers at the times the drugs were applied and tell us whether and how the tether lengths changed before and after drug addition. The differences in tether lengths between start and finish of backward movement is the total distance traveled backward by partner half-bivalents towards each other, which in turn was recorded as the fractional distance of the initial tether that the chromosomes moved backward. Slopes of movement *versus* time on the graphs (the lines are computer generated lines of best fit) tell us movement velocities.

**Figure 2A.**
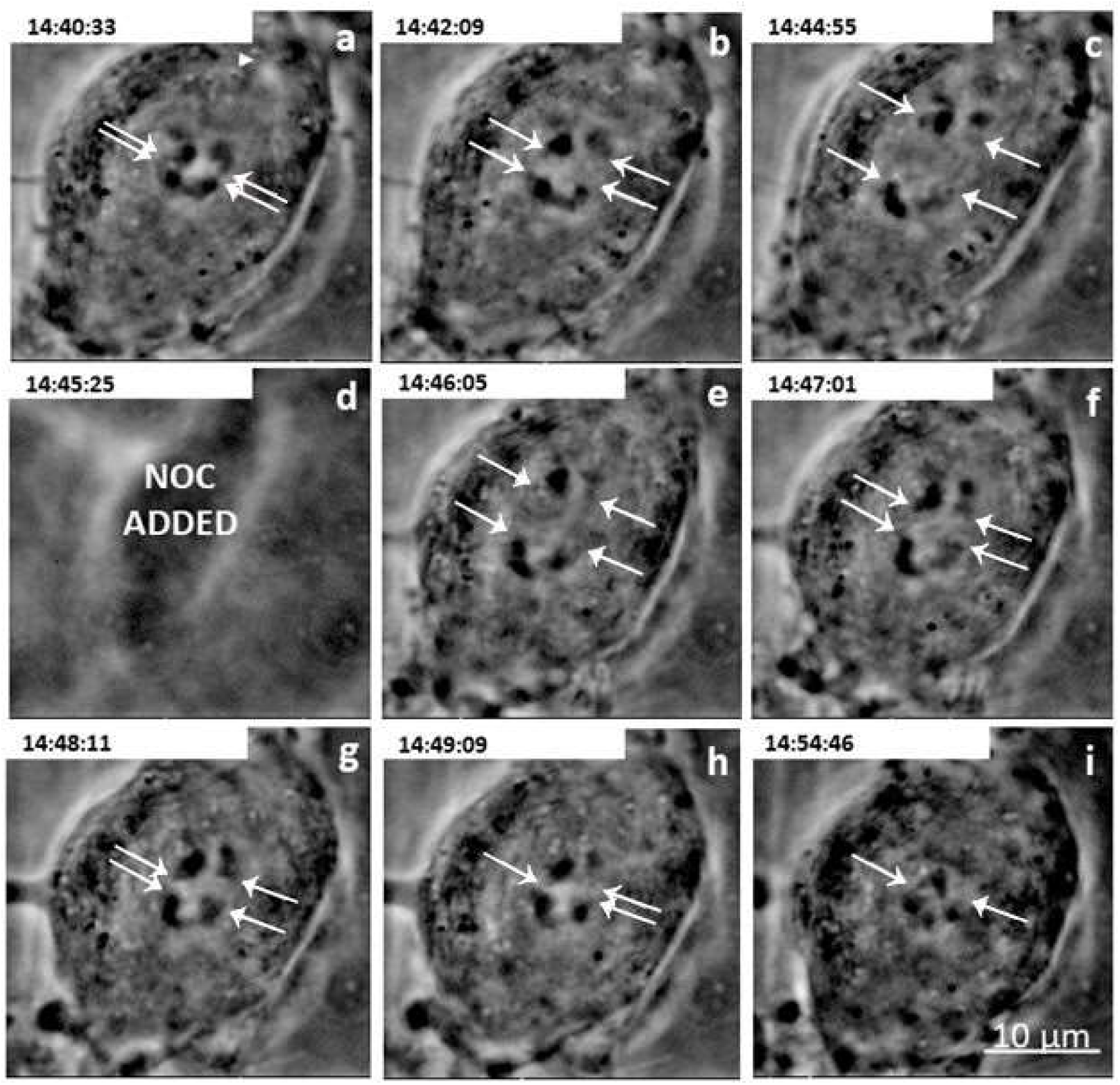
A cell treated with **30µM Nocodazole**. Times (hrs: min:sec) are at the top of each panel. **(a)** In this plane of focus two pairs of chromosomes start anaphase; telomeres of partner half-bivalents are indicated by arrows, here and in subsequent images. The arrowhead indicates the point near the top spindle pole used as a fixed reference point for measurements. **(b-c)** Partner half-bivalents continue separating. **(d)** Nocodazole (NOC) was added. **(e)**The half-bivalents stop separating and start moving backwards towards each other. **(f-h)** partners continue moving backwards towards each other. **(i)** Backward movement stops as partner telomeres contact each other. Scale bar in (i) represents **10 µm.**

**Figure 2B.**
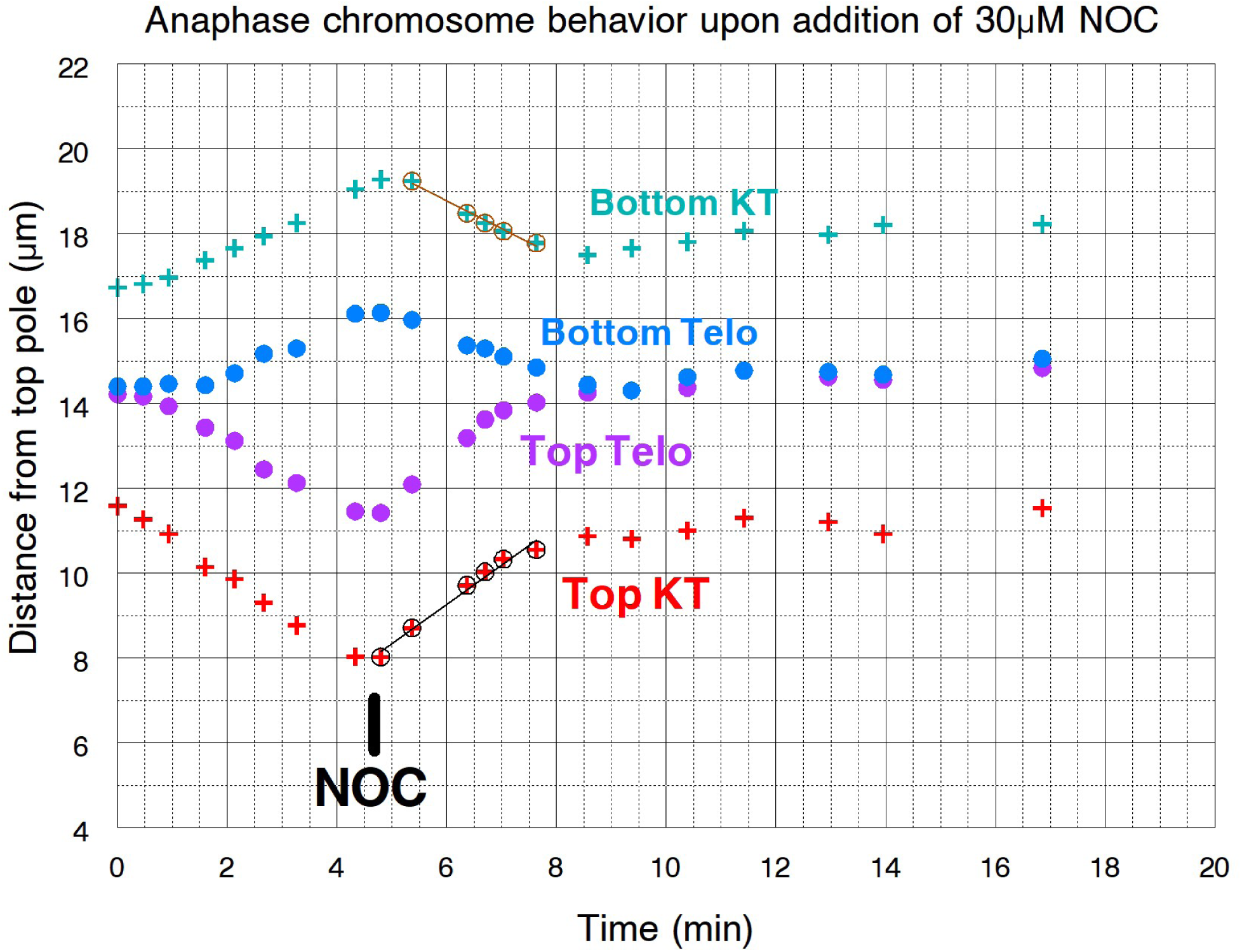
Movement graph for the left pair of half-bivalents illustrated in Figure 2A. Time = 0 min on the graph corresponds to 14:40:33 (***panel a***) in Fig. 2A. Partner *kinetochores* are indicated as Top KT and Bottom KT, partner *telomeres* as Top Telo and Bottom Telo. Partners were moving apart when Nocodazole (NOC) was added (indicated on the graph as NOC) after which they stopped moving poleward and moved backward until they met each other. Slopes of the lines represent movement velocities.

**Figure 3A.**
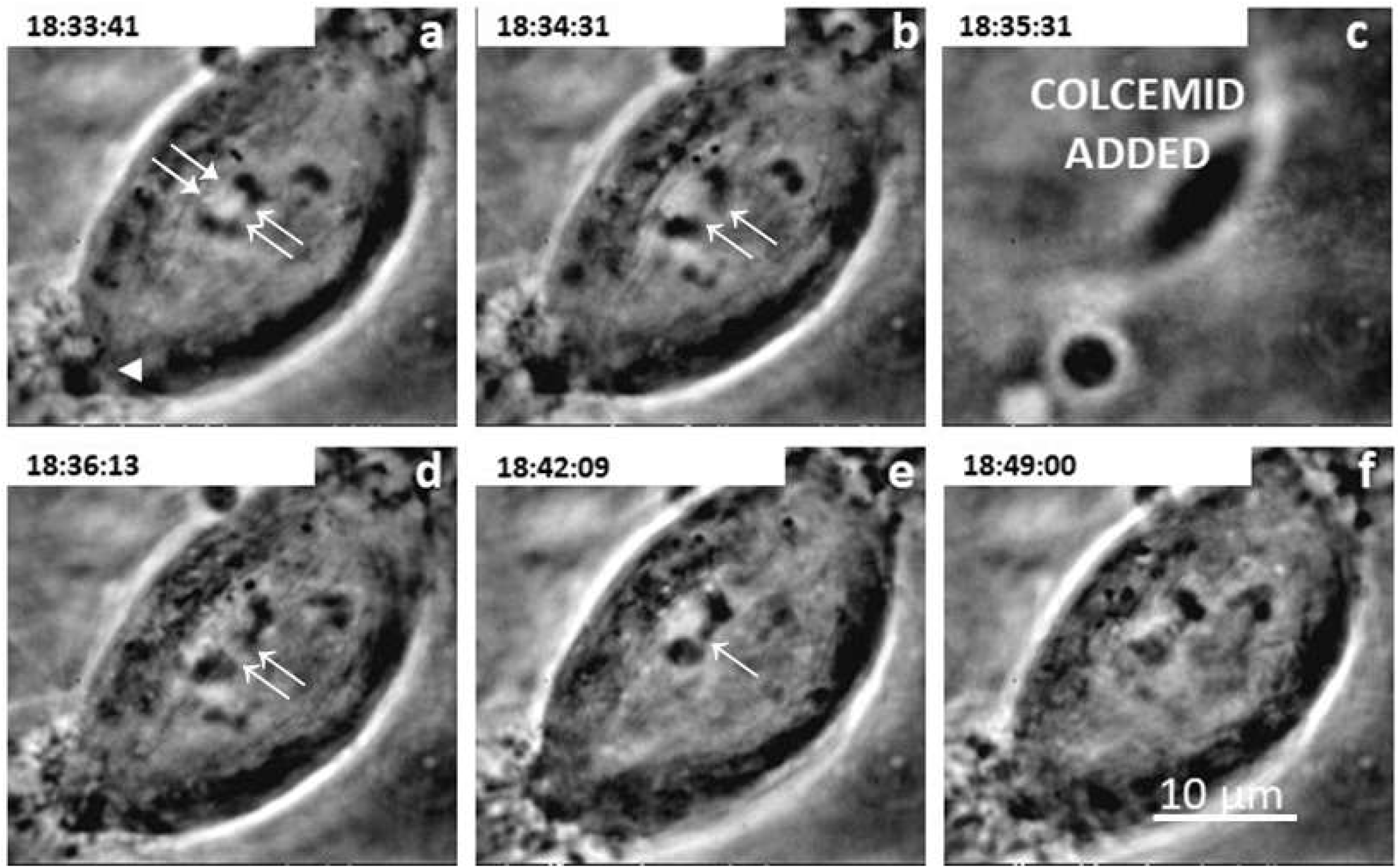
A cell treated with **100µM Colcemid.** Time (hrs: min:sec) is indicated at the top of each panel. **(a)** Arrows point to the partners of two pairs of chromosomes starting anaphase separation. The arrowhead points to the point near the bottom spindle pole used as a fixed reference point for measurements. **(b)** Partner half-bivalents continue separating. **(c)** Colcemid added. **(d)** Half-bivalents stop separating and start moving backwards towards each other. **(e)** Backward movement stops as partner telomeres meet. **(f)** Partners remain as they were. Scale bar in (f) represents **10 µm.**

**Figure 3B.**
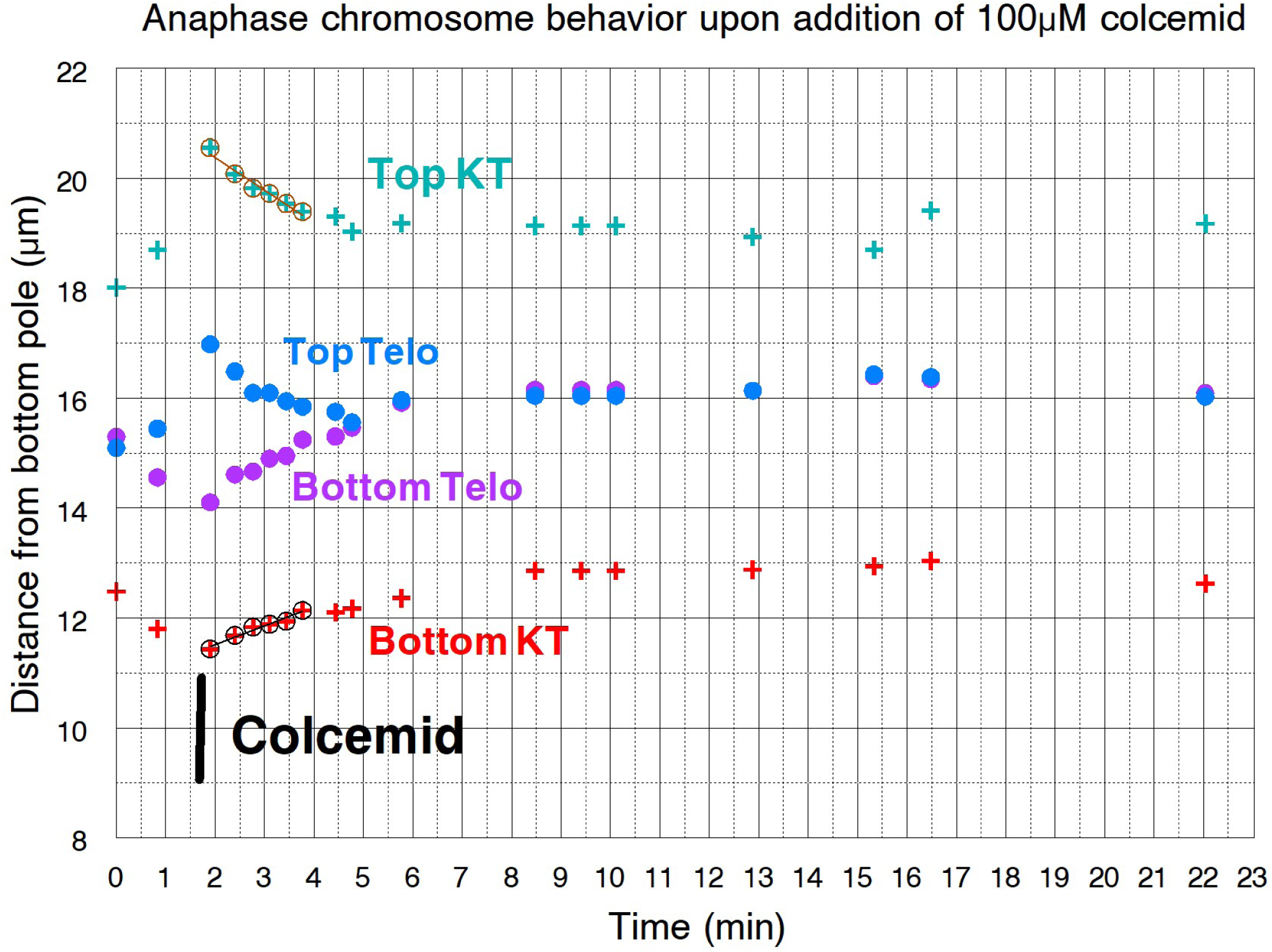
Movement graph for the left pair of chromosomes illustrated in Figure 3A. The graph time=0 min corresponds to 18:33:41 (***panel a***) in Fig. 3A. Partner *kinetochores* are marked as Top KT and Bottom KT, partner *telomeres* as Top Telo and Bottom Telo. Partners were moving apart when Colcemid was added after which they stopped moving poleward and moved backward until they met. Slopes of the lines represent velocities.

**Figure 4A.**
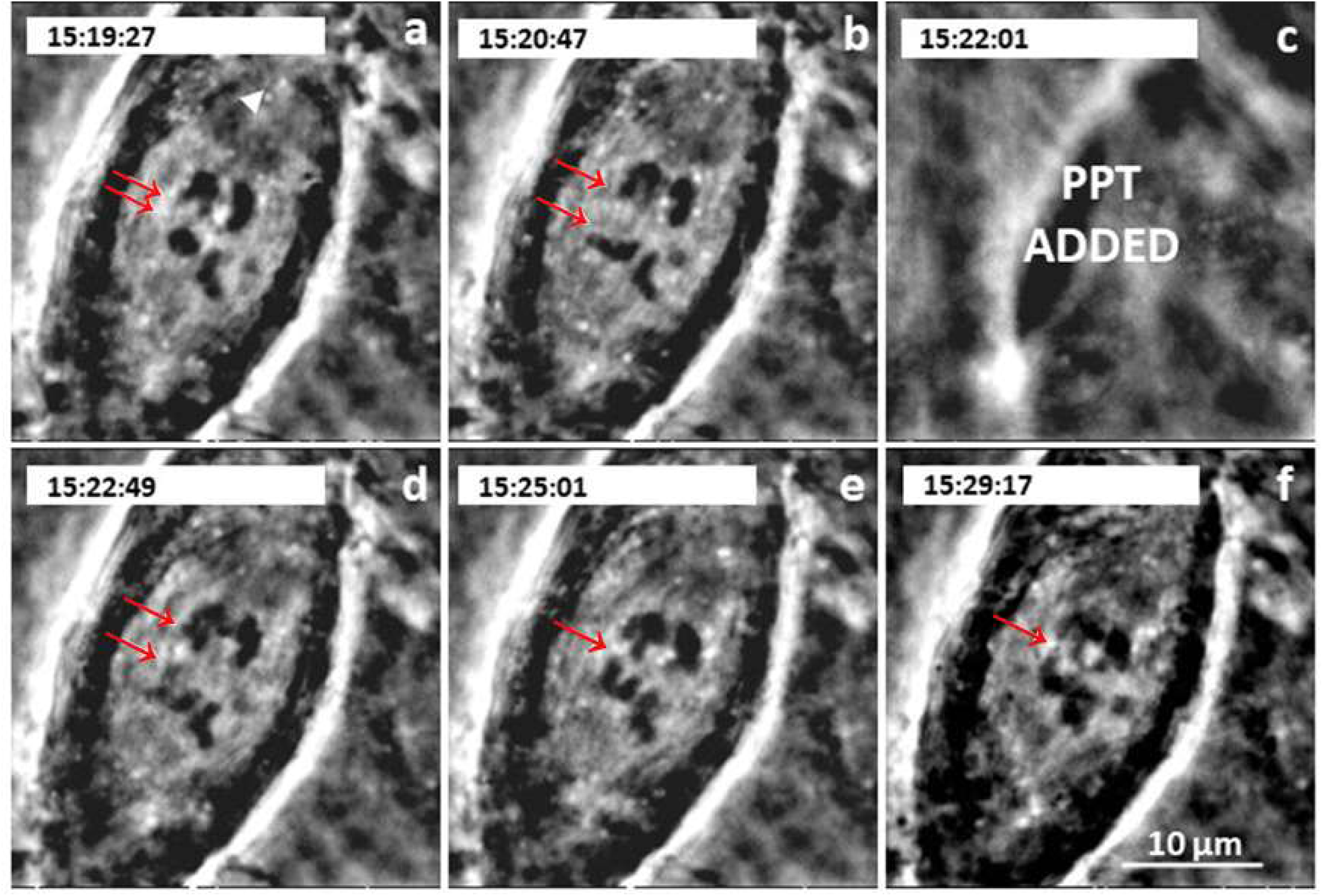
Cell treated with **60µM Podophyllotoxin**. Time (hrs: min:sec) is indicated at the top of each panel. **(a)** Arrows point to the partner telomeres of one pair of half-bivalents; both visible pairs are starting anaphase separation. The arrowhead points to the point near the top spindle pole used as a fixed reference point for measurements. **(b)** Partner half-bivalents continue separating. **(c)** 60µM Podophyllotoxin (PPT) added. **(d)** Half-bivalents stop separating and start moving backwards towards each other. **(e)** Backward movement continues. **(f)** Backward movement stops as partner telomeres meet, then remain as they are. Scale bar in **(f)** represents **10 µm.**

**Figure 4B.**
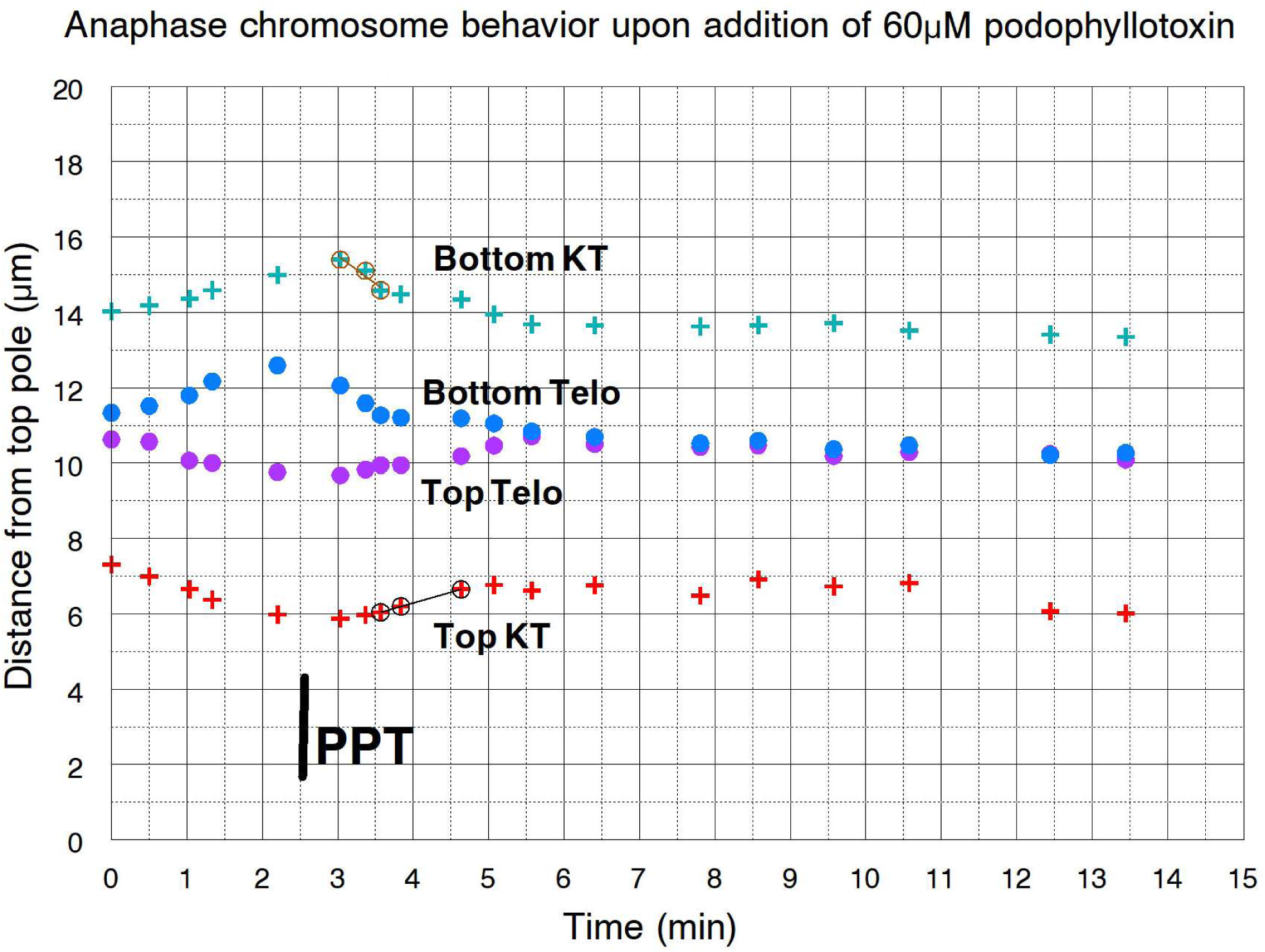
Movement graph for the left pair of homologous chromosomes illustrated in Figure 4A. The graph time = 0 corresponds to 15:19:27 in Figure 4A. Partner *kinetochores* are marked as Top KT and Bottom KT, partner *telomeres* as Top Telo and Bottom Telo. Partners were moving apart when podophyllotoxin (PPT) was added, after which they stopped moving poleward and moved backwards until the telomeres met. Slopes of the lines represent velocities.

Chromosomal behavior was similar for each of the microtubule depolymerizing agents, nocodazole, colcemid, and podophyllotoxin,. Poleward movement stopped very soon after the drugs were added. Very shortly thereafter the chromosomes moved backward until the telomeres met, or until the telomeres stopped moving backward without meeting Since all three drugs depolymerize microtubules, albeit via different mechanisms, and all had the same effects, we grouped the results together. Overall, 107 homologous pairs of partner half-bivalents were observed in 38 cells treated with the three inhibitors; all inhibitors stopped anaphase chromosomal segregation within 1 minute of drug treatment (Table 1). If the backward movements are due to mitotic tethers, since tethers become less elastic as anaphase proceeds one would expect that whether or not the chromosomes move backward, and how far they move, depends on the tether length at the time the drug was added.

**Table 1.** Anaphase segregation arrest in cells treated with microtubule depolymerizing drugs. PPT = Podophyllotoxin

| **Drug Treatment** | **Treated cells** | **Half-bivalent pairs that stopped segregating/**<br>**number of half-bivalent pairs observed** |
| --- | --- | --- |
| 20 μM NOC | 3 | 9/9 |
| 30 μM NOC | 17 | 48/48 |
| 60 μM NOC | 4 | 10/10 |
| 90 μM NOC | 1 | 3/3 |
| 100 μM Colcemid | 2 | 6/6 |
| 200 μM Colcemid | 2 | 6/6 |
| 10 μM PPT | 1 | 3/3 |
| 20 μM PPT | 1 | 3/3 |
| 30 μM PPT | 1 | 3/3 |
| 50 μM PPT | 3 | 8/8 |
| 60 μM PPT | 3 | 8/8 |
| **TOTALS** |  | **107/107** |

#### After treatment of cells with microtubule depolymerizing drugs, *shorter tethers* are more consistently elastic and cause backward chromosomal movement more frequently than *longer tethers*

To determine whether the backwards movements were due to tethers we tested whether there was a correlation between frequency of backward movement and the tether length when depolymerizing agents were added. In experiments when arm fragments are formed by cutting arms with a laser, shorter tethers are more consistently elastic than longer tethers and shorter tethers cause backward chromosomal movements more often than longer tethers (LaFountain et al., 2002; Forer *et al*., 2021). The same should apply when microtubule depolymerizing agents are added if the backwards chromosome movements are due to mitotic tethers. We measured backward movements as a function of tether lengths at the time of drug addition (Figure 5). All the backward movements started within 1 minute of anaphase arrest, and, as with arm fragments (Forer et al., 2021), whether chromosomes moved backward or not depended on tether length at the time the depolymerizing agents were added: when the drugs were added at short tether lengths (<3µm) chromosomes moved backward 100% of the time, fewer moved backwards at longer lengths, and none moved backwards at tether lengths ≥9 µm (Figure 5). This pattern of decreasing frequencies with increasing tether length is characteristic of arm fragment movements (La Fountain et al., 2002; Forer et al., 2021), as seen in Figure 7C in (Forer et al., 2021), and is consistent with the idea that the chromosomes move backward after treatment with microtubule depolymerizing agents because of forces from tethers.

**FIGURE 5.**
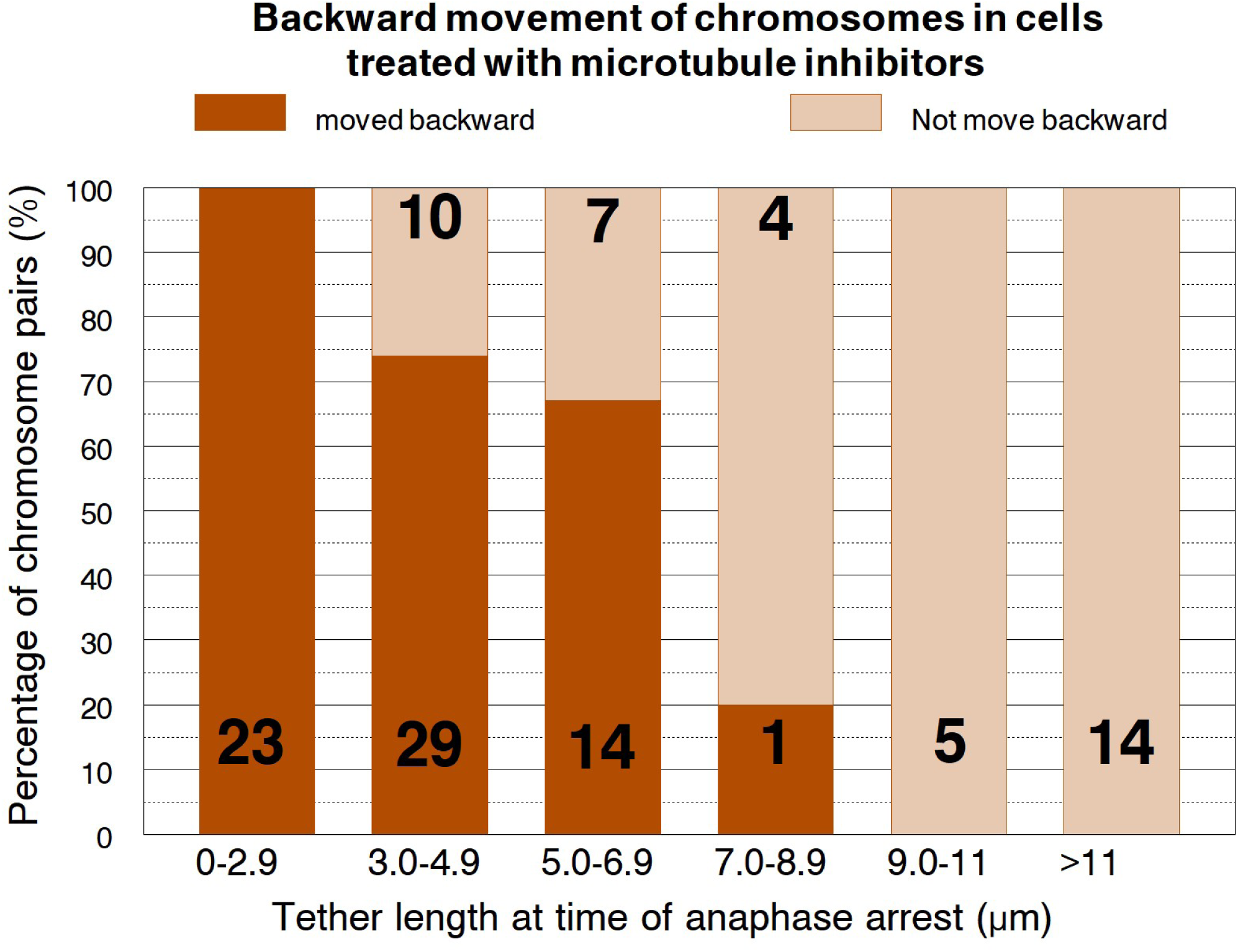
Backward movement of half-bivalents in cells treated with microtubule depolymerising agents at different tether length. No half-bivalents moved backwards at tether lengths above 9µm; 100% moved backward at tether lengths less than 3µm. The results are similar to results on arm fragments (from laser experiments) shown in Graph 7C of Forer et al. (2021)

#### Shorter tethers cause backward chromosomal movements *over proportionally longer distances* than longer tethers do after cells are treated with microtubule depolymerization agents

We evaluated tether elasticity after treatment with microtubule depolymerizing drugs because when arm fragments move, shorter tethers tend to shorten more in proportion to their initial length than longer tethers do (Forer et al., 2021). The extent of tether shortening was calculated as the distance the telomere in question moved backward as a fraction of the initial tether length, as shown in (Figures 6A, 6B). In those figures we grouped the results from all three microtubule depolymerising agents that we used. Both graphs clearly show that shorter tethers cause backward movements over greater fractional distances of the tethers than longer tethers do. The same results occur in arm fragments formed at different length tethers in untreated (control) cells (e.g., Figures 6A and 6B in Forer *et al.,* 2021), buttressing the argument that mitotic tethers cause the backward movements of chromosomes after anaphase cells are treated with microtubule depolymerizing agents. Further statistical analysis confirms this conclusion.

**Figure 6A.**
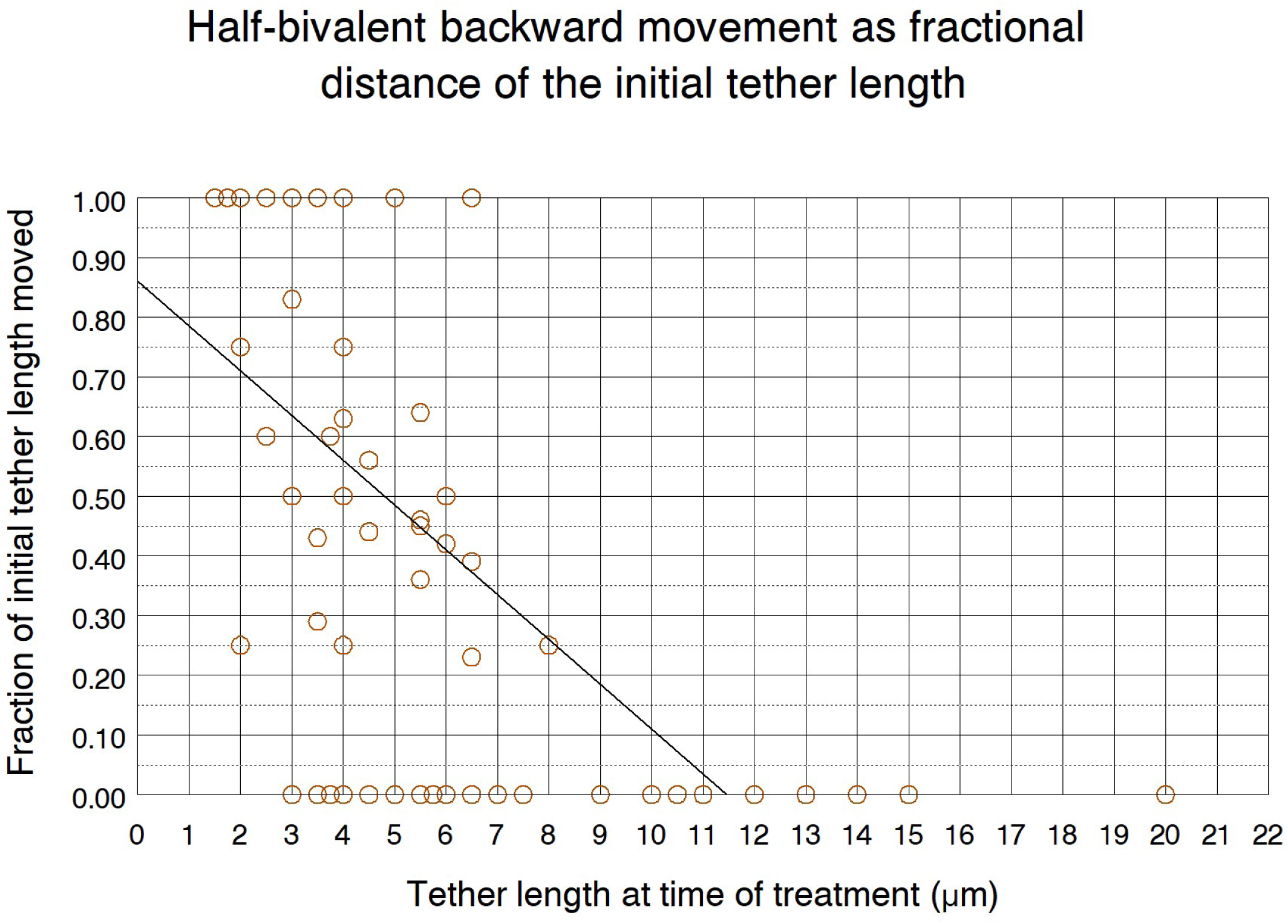
Cells treated in anaphase. The scatter diagram indicates distances moved backwards by half-bivalent pairs as fractions of their initial tether lengths. The line was obtained using Spearman’s rank correlation. The correlation coefficient (r_s_) was -0.724, and with alpha level ≤ 0.01, the p value equaled 1.34x10^-18^ . Thus chromosomes move backward in our experiments with characteristics similar to arm fragments severed from arms (Forer et al., 2021).

**Figure 6B:**
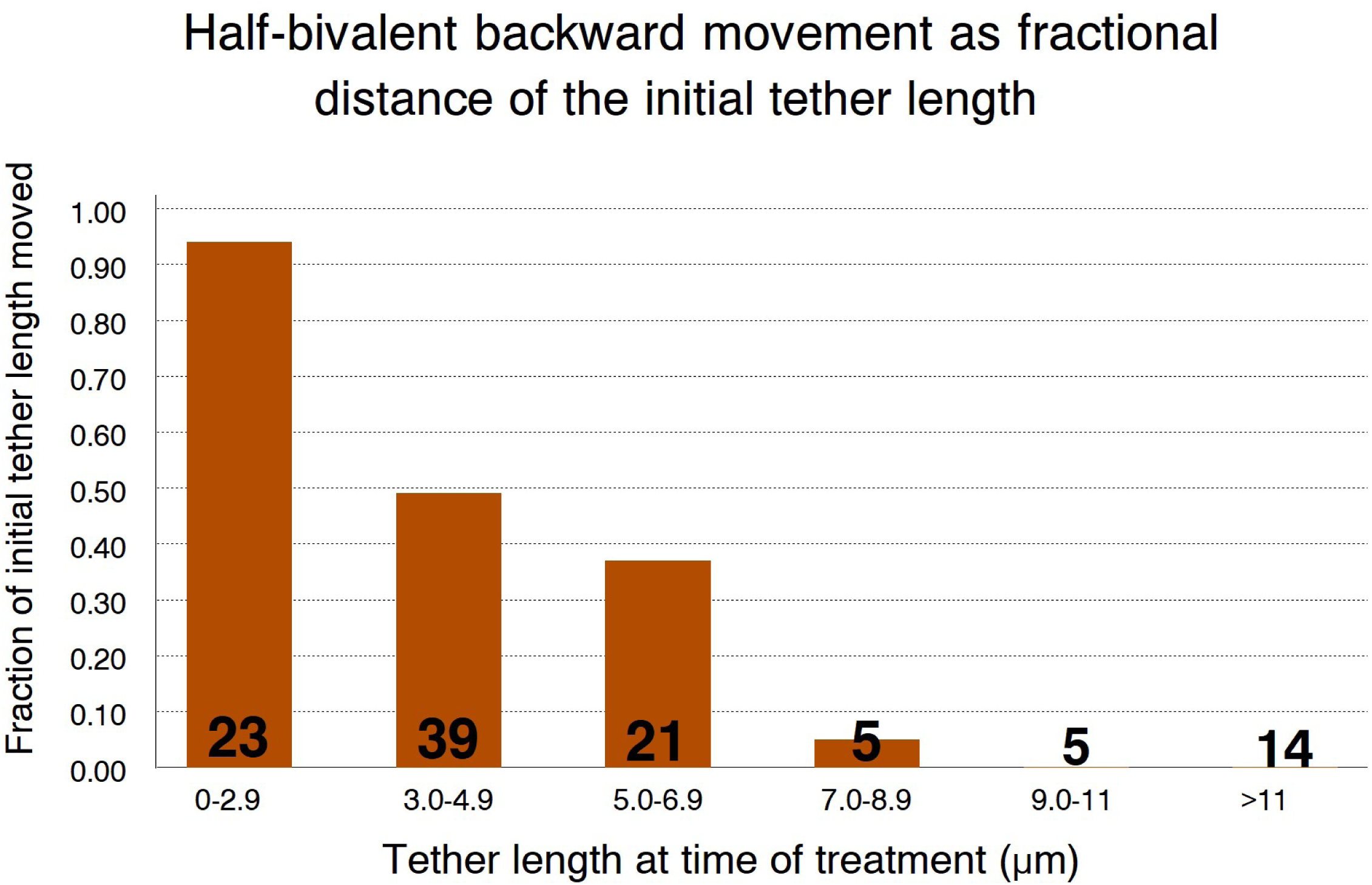
a bar graph of the scatter plot data from Figure 6A. The values were grouped into tether length ranges; average fractional distances moved backwards in each category were plotted for each tether-length range. 107 bivalent pairs were examined in total. Numbers in each tether length category are written on the bars. The results are similar to graphs of tether lengths *versus* fraction of tether lengths that arm fragments moved [e.g., Figure 6B in Forer et al., 2021].

Spearman’s rank correlation was calculated to verify that shorter tether elasticity was unlike longer tether elasticity, and that the differences between the fractional backward chromosomal movements caused by tethers of different lengths were statistically significant. The correlation coefficient (rs) was calculated as -0.724; with alpha level ≤ 0.01, p value equals 1.34x10^-18^. Therefore, when cells are treated with compounds that depolymerise microtubules, the fractional distances moved backward by chromosomes due to their tether elasticity is negatively correlated to their tether length more than 99.99% of the times (Figures 6A, 6B). This highly significant correlation further corroborates that the backwards movement we observe are due to tethers, and therefore that tethers function to move half-bivalents backwards after cells are treated with microtubule depolymerizing agents as they do to move arm fragments backwards.

#### The velocities of backward chromosomal movements at different tether lengths are statistically the same after cells are treated with microtubule depolymerising drugs

Chromosomal arm fragments cut at different tether lengths move backward with statistically identical velocities in both CalA-treated and untreated cells (Forer *et al*., 2021). Similarly, in cells treated with dilute lysis buffer, half-bivalents with different tether lengths move backward with velocities that are statistically the same (Aidil and Forer, 2024). The same holds true in the present study: Figure 7 illustrates the backward chromosomal velocities grouped together from all treatments. We conclude that the microtubule depolymerising agents used herein do not adversely affect tether function relative to how they function to move arm fragments and therefore the various microtubule inhibitors used in this study do not alter this characteristic of tether behaviour.

**Figure 7.**
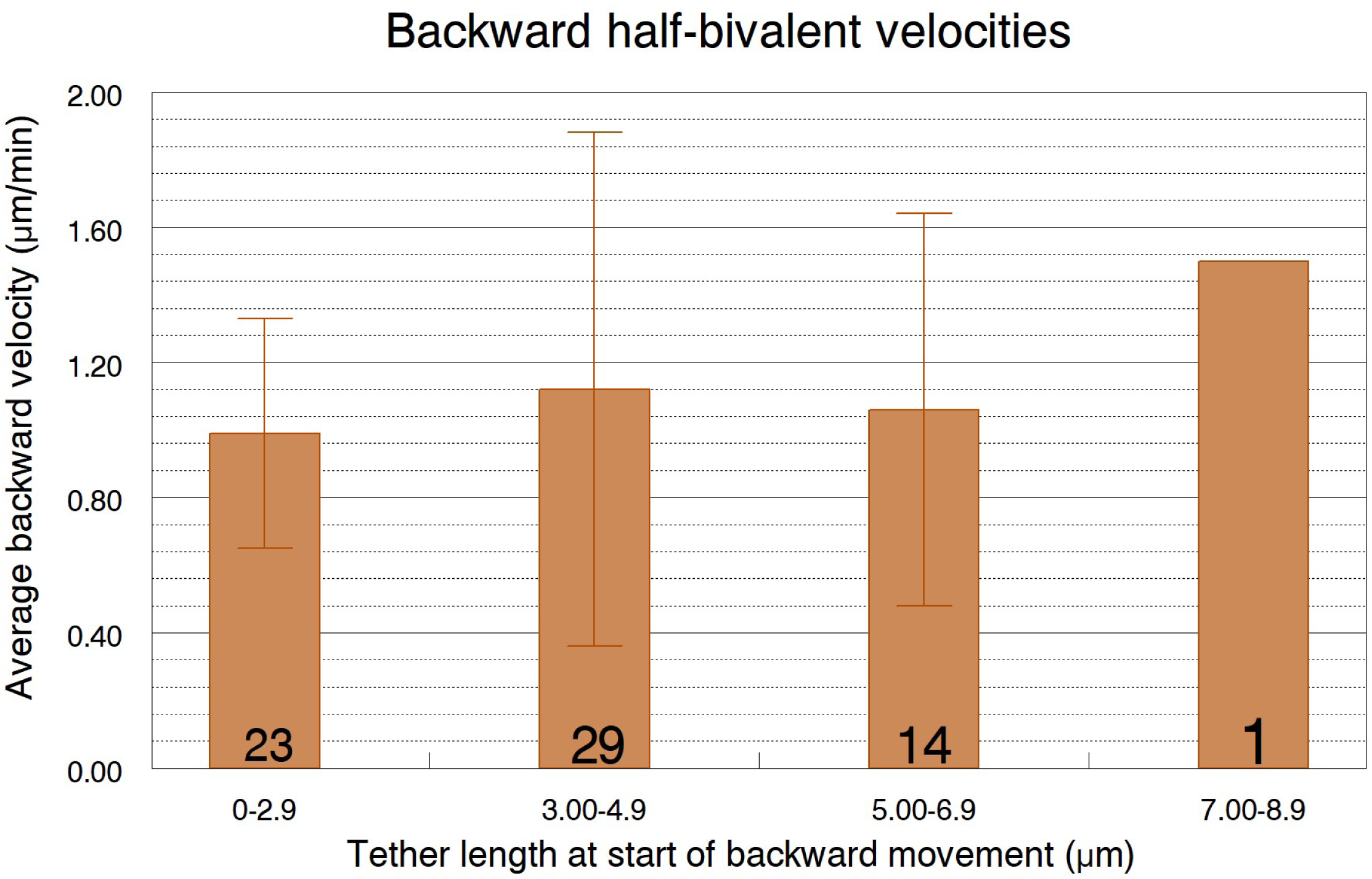
Average backward half-bivalent velocities versus tether lengths after treatments with microtubule depolymerizing agents. Standard deviations are indicated on the bars. N = 67 half-bivalent pairs; the numbers within each grouping are indicated on the respective bars. Student’s t-test was performed on all data sets and all differences are statistically insignificant (alpha level set at 0.01).

#### Backward chromosomal velocities after addition of microtubule deoplymerising agents are statistically similar to backward chromosomal velocities seen with other anaphase arresting treatments

Elastic tethers move chromosomes backward during anaphase arrest in microtubule-inhibited cells as they do during anaphase arrest in partially lysed cells (treated with dilute lysis buffer), and as they do in CalA-treated partially lysed cells (treated with CalA then lysis+ CalA) (Adil and Forer, 2024). How do the backward movement velocities in our study compare with those in other treament conditions? Average backward velocities versus tether lengths were compared and are statistically the same using Student’s t-test (Figure 8), except perhaps for comparison of microtuble-inhibition and dilute-lysis treatment groups at tether lengths <5 µm: the t-test p-value was 0.011, meaning that there was a 0.011 probability of the two groups being different. The t-test probability for all other group pairings was >0.01, above our acceptance level, so backward chromosomal velocities seen with all these anaphase arresting treatments are statistically the same.

**Figure 8.**
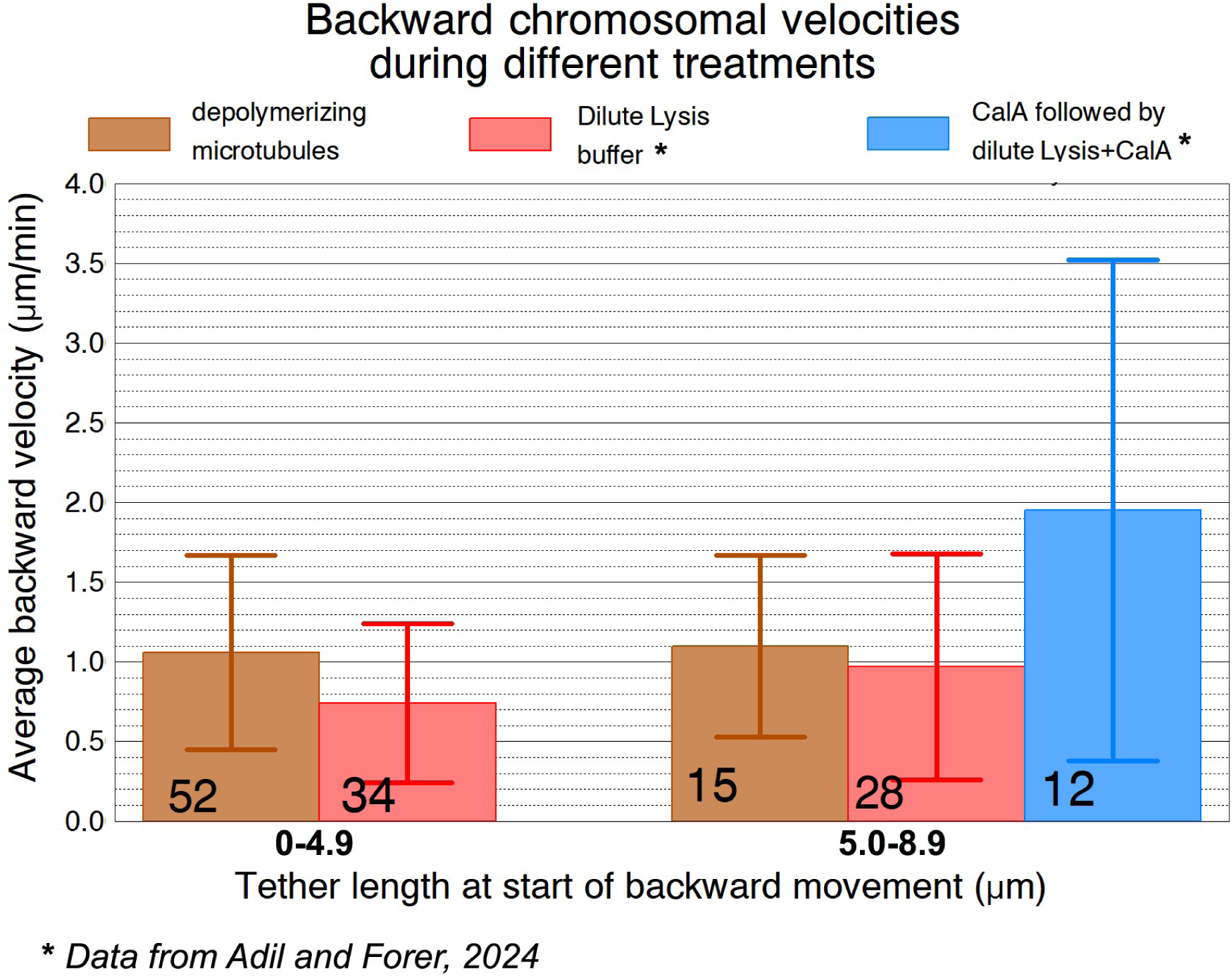
Average backward velocities with their standard deviations (capped bars) shown at different tether length ranges during treatment with microtubule depolymerizing agents. The data on depoly,merizing microtubules are from this article. Other data, from Adil and Forer (2024), are for treatment with dilute lysis buffer, and for treatment first with CalA and then with dilute lysis buffer in the continued presence of CalA. The numbers in each category are written on the respective bars. Student’s t-test was performed on all data sets and the differences were statistically insignificant (alpha level set at 0.01) except the two values <5.0µm, which had an alpha level of 0.011, meaning a 0.011 probability of being different.

#### None of the depolymerizing agents depolymerized acetylated kinetochore microtubules

We used confocal microscopy to determine the status of spindle microtubules after treatment with depolymerizing agents. Because chromosomes moved at speeeds of 1-1.5 µm/min (Figure 7), much slower than the (4-7 µm/min) backward speeds of chromosomal arm fragments (Forer *et al*., 2021), we suspected that the microtubule depolymeriszing drugs did not disassemble kinetochore-microtubules, and that the attached kinetochore microtubules slowed the backward movements. We tested this supposition by immunostaining nocodazole-treated cells for total tubulin and for acetylated tubulin. We stained control cells and we stained cells treated for 25 minutes with 30 µM nocodazole. In *control* cells there was staining of both non-kinetochore microtubules and kinetochore microtubules (Figure 9A). The only staining for acetylated microtubules was in the kinetochore microtubules (Figure 9B), as shown previously for crane-fly spermatocytes by Wilson and Forer (1989) and Wilson et al. (1994). In *cells treated with nocodazole*, and stained for total tubulin, there were no non-kinetochore microtubules: the acetylated tubulin and total tubulin images looked identical (Figure 9D and 9E). It appears that the non-kinetochore microtubules were depolymerized by the microtubule depolymerizing drugs but the acetylated kinetochore microtubules were not. We suggest that these remaining kinetochore microtubules most likely resist and slow down the backward movements of the chromosomes.

**FIGURE 9.**
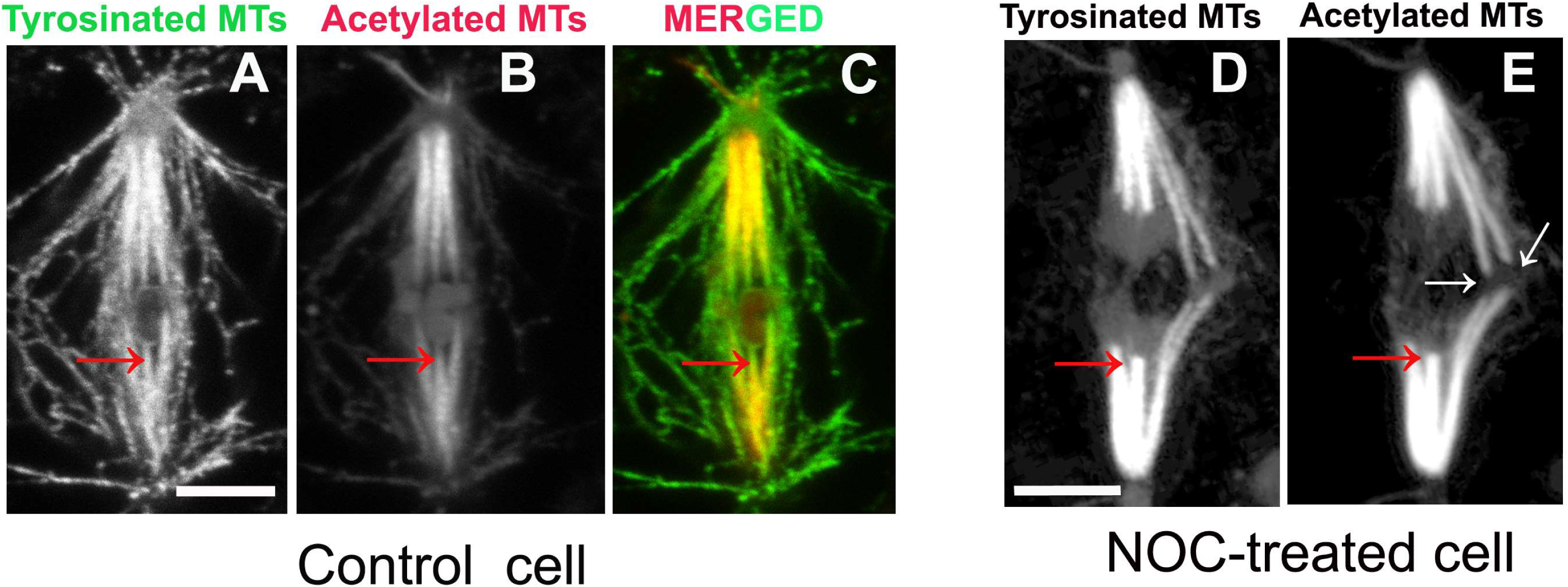
Staining for tyrosinated (total) tubulin and acetylated tubulin, as indicated. ***Figs 9A to C*** illustrate one ***control cell*** just starting anaphase. The acetylated tubulin stain is red and stains only the kinetochore microtubules and the two flagella at each pole. One kinetochore fibre microtubule bundle is indicated by a red arrow. The tyrosinated tubulin stain is green, and stains both non-kinetochore microtubules and kinetochore microtubules. The kinetochore microtubules are not acetylated near the kinetochore, as seen in Fig. **9A** and **9B** (compare the kinetochore fibre pointed to be the red arrows), As illustrated in the merged colour image in Fig. **9C**, there is no acetylated microtubule staining near the kinetochore so the microtubules near the kinetochore stain green (tyrosinated) and the kinetochore microtubules only stain for acetylated tubulin closer to the pole. The uneven staining is because the acetylation occurs primarily in microtubules, not in tubulin monomers; acetylation takes time, and after the monomers enter the kinetochore microtubules at the kinetochore they move immediately to the poles (e.g., LaFountain et al., 2004). They become acetylated closer to the poles only after they have been stable for some minutes (e.g., Wilson and Forer,1997). ***Figs. 9D and E*** illustrate one cell that was ***treated with 30µM nocodazole*** in anaphase. Only kinetochore microtubules are present. One kinetochore fibre is indicated by a red arrow. The treatment with Nocodazole seems to have stopped incorporation of new tubulin, because after nocodazole treatment there is no gap in staining of acetylated tubulin near the kinetochore. The white arrows point to the two sex chromosomes at the equator, which seem to have been pushed out of the spindle, perhaps because nocodazole causes spindle shortening. The scale bars (in ***Figs 9A*** and ***9D***) represent 5µm. The image of the control cell is the sum of 6 superposed confocal slices: the image of the nocodazole-treated cell is the sum of 13 slices.

## DISCUSSION

The major finding of this research is that treatments of cells with microtubule depolymerizing agents do not alter the functioning of mitotic tethers: the tethers still exert anti- polar forces on telomeres in the same manner they do in untreated living cells. The same holds true when cells are partially lysed (Aidil and Forer, 2024) and is strong indication that the action of mitotic tethers in producing backward force on separating chromosomes does not require functioning of the usual mitotic forces. Tethers not only function in the absence of functional microtubules, they function with the same properties as movements of arm fragments severed from chromosome arms during anaphase (Forer et al., 2021). Tether properties after anaphase arrest in partially lysed cells are the same as after anaphase arrest from microtubule depolymerizing drugs, and the same as in movements of arm fragments formed in anaphase, namely: shorter tethers cause more frequent chromosomal backward movements than longer tethers (Figure 5); shorter tethers cause backward movements over greater fractional distances (of the tether) than longer tethers (Figure 6); and tethers of different lengths cause backward movements with statistically the same velocities (Figures 7, 8). Thus, half-bivalent movements in these systems are identical to those of arm fragments (Forer et al., 2021), with *one exception*, that the speeds of the arm fragments are considerably faster than the speeds of backwards movements of whole chromosomes in cells, including the speed of backward movement after adding Calyculin A in early anaphase (Fabian et al., 2007). We think there is a simple reason why chromosome arm fragments move faster: chromosomes are encumbered by attached kinetochore microtubules whereas chromosome arm fragments are not. As we have shown, the acetylated microtubules are not depolymerized and kinetochore microtubules remain attached to the chromosomes (Figure 9).

In sum, all characteristics of tethers deduced from cutting arms and observing the movements of the resultant arm fragments are reproduced when the mitotic apparatus is treated with microtubule depolymerizing agents, except for the speed that the chromosomes move backward. We suggest that the attached kinetochore microtubules slow the backwards movements.

Our results have implication to two other issues. One is the study of mitotic tethers. Most previous studies of tethers involved the use of a laser microbeam to cut arms from anaphase chromosomes: there was no other way to study tethers. As the results of this study and several previous studies (Kite and Forer, 2020; Adil and Forer, 2024) show, one can study tethers using straightforward light microscopy, as we have done, without need for a specialized laser set-up. There is one possible experimental qualification, though. In crane-fly spermatocytes the half-bivalents move to a stationary people: anaphase B, spindle pole elongation, does not occur until the autosomes reach the poles. Thus if anaphase chromosome separation is due primarily to spindle elongation, the functional tethers may not cause the chromosomes to respond in the same way as in crane-fly spermatocytes because in other cells the chromosomes might be connected to the elongating spindle.

The other issue is how the microtubule depolymerisers block anaphase chromosome movements. During anaphase chromosomal poleward movement in crane-fly spermatocytes, as the attached chromosomes move towards their poles new tubulin monomers add to the kinetochore microtubules at the kinetochore simultaneously with tubulin monomers being removed from the kinetochore microtubules at the poles (LaFountain *et al*., 2004). The subunits at the poles are removed from the kinetochore microtubules at a faster rate than they are added at the kinetochore, which causes the poleward movement (LaFountain *et al.,* 2004). The depolymerizing agents Nocodazole, colcemid, and podophyllotoxin block microtubule polymerization (Bryan, 1974; LaFountain, 1985; Jordan *et al*., 1992; Silverman-Gavrila and Forer, 2000; Hamel, 2003). Blocking polymerisation means that new tubulin does not enter kinetochore microtubules at the kinetochore, thereby causing the then stationary tubulin subunits at the kinetochore to become acetylated (Figure 9; Wilson et al., 1994; Wilson and Forer, 1997). The treatments also block subunits from leaving microtubules at the pole, and thereby halt anaphase chromosome movements, after which the mitotic tethers pull the half- bivalents backward.

In conclusion, this research has shown that microtubule depolymerization drugs arrest anaphase chromosomal poleward movements. Inhibiting chromosome movement to the poles does not affect the functioning of mitotic tethers, so the tethers cause the chromosomes to move backward. Because the chromosomes are held back by the attached kinetochore microtubules, the chromosomes move backward slower than arm fragments move when they are cut from chromosome arms in non-treated cells.

## ACKNOWLEDGEMENTS

Some of the data in this article was part of a M Sc thesis submitted by AA to York University in 2023. This work was supported by grant RGPIN-2019-06299 from the Natural Sciences and Engineering Research Council of Canada (NSERC) to AF. During part of this work AA was recipient of a Natural Sciences and Engineering Research Council of Canada Graduate Scholarship-Master’s program (NSERC-CGS M) award.

